# An atlas of plant full-length RNA reveals tissue-specific and evolutionarily-conserved regulation of poly(A) tail length

**DOI:** 10.1101/2022.01.21.477033

**Authors:** Jinbu Jia, Wenqin Lu, Bo Liu, Yiming Yu, Xianhao Jin, Yi Shu, Yanping Long, Jixian Zhai

## Abstract

Poly(A) tail is a hallmark of eukaryotic mRNA and its length plays an essential role in regulating mRNA metabolism^1–3^. However, a comprehensive resource for plant poly(A) tail length has yet to be established. Here, we applied a poly(A)-enrichment-free, Nanopore-based method^4,5^ to profile full-length RNA with poly(A) tail information in plants. Our atlas contains over 120 million polyadenylated mRNA molecules from seven different tissues of Arabidopsis, as well as the shoot tissue of maize, soybean and rice. In most tissues, the size of plant poly(A) tails shows peaks at approximately 20 and 45 nt, presumably the sizes protected by one and two poly(A) binding proteins (PABP), respectively^2,6^, while the poly(A) tails in pollen exhibit a distinct pattern with strong peaks centered at 55 and 80 nt. Moreover, poly(A) tail length is regulated in a gene-specific manner — mRNAs with short half-lives in general have long poly(A) tails, while mRNAs with long half-lives are featured with relatively short poly(A) tails that peak at ~45 nt, suggesting that protection of poly(A) tail in this size by PABP is essential for mRNA stability. Across species, poly(A) tails in the nucleus are almost twice as long as in the cytoplasm, implying a conserved rapid shortening process of poly(A) tail occurs before the mRNA is stabilized in cytoplasm. Our comprehensive dataset lays the groundwork for future functional and evolutionary studies on poly(A) tail length regulation in plants.

## Main text

A non-template poly(A) tail is the hallmark of eukaryotic mRNA^1–3^ and is vital in promoting translation and protecting mRNAs integrity with the help of cytoplasmic poly(A) binding proteins (PABPC)^2,7,8^. The length of the poly(A) tail is dynamically regulated by poly(A) polymerase and deadenylase, and shorting poly(A) tails to a certain threshold would release PABPC and trigger mRNA decay^2,3,9–12^. A growing body of evidence reveals that altering poly(A) tail length plays important role in regulating gene expression^2,9,13,14^. In the last decade, a few high-throughput Illumina-based methods have been developed for genome-wide characterization of poly(A) tail length, including PAL-seq^13^, TAIL-seq^15^, mTAIL-seq^16^, PAT-seq^17^ and TED-seq^18^. Advances in long-read sequencing platforms such as PacBio and Nanopore have also enabled the development of methods that detect full-length mRNA with poly(A) tail information, such as FLAM-seq^19^, PAIso-seq^20^, Nanopore direct RNA sequencing (DRS)^21–23^, and one developed by our group named FLEP-seq (Full-Length Elongating and Polyadenylated RNA sequencing)^4,5^. However, existing resources on plant poly(A) tails are still limited^8,13,21,24–26^. Therefore, establishing a comprehensive landscape for poly(A) tail length from various tissue types and in different species would greatly facilitate the study of poly(A) regulation in plants.

Here, for low-input and cost-efficient detection of poly(A) tails, we further optimized FLEP-seq procedure for studying total RNA and streamlined library construction with the newly released Nanopore PCR-cDNA sequencing kit which uses ligase-free attachment of rapid sequencing adapter to bypass the steps of ds-cDNA repair, A-tailing and adapter ligation — we named this new version of the protocol as FLEP-seq2 (Fig. 1a and Fig. S1). Because FLEP-seq is originally designed to detect nascent RNAs that may or may not have a poly(A) tail, it does not involve any step for selecting poly(A) tail. This turns out to be a unique advantage for unbiased measurement of poly(A) tail length, as all other long-read methods (FLAM-seq, PAIso-seq, and DRS) use oligo-dT to select poly(A)+ mRNAs, which could disfavor mRNAs with a short poly(A) tail (Fig. 1a). Indeed, compared to published DRS data of the same tissue^21^, FLEP-seq2 detects more transcripts with short poly(A) tail, and the poly(A) length distribution of transcripts shows the highest peak at ~20 nucleotides (nt) (Fig. 1b, upper panel). Despite this slight difference, the overall poly(A) length distributions obtained by FLEP-seq2 and DRS are highly similar (Fig. 1b, upper panel). The median poly(A) length of genes between FLEP-seq2 and DRS (*r*=0.72) (Fig. 1b, lower panel, Fig. S2a), as well as between the two biological replicates of FLEP-seq2 (*r*=0.87) (Fig. S2b), are highly consistent. In addition, FLEP-seq2 has a significant advantage in term of output per Nanopore flow cell compared to DRS — on a regular Nanopore MinION flow cell, one FLEP-seq2 library can yield ~10-20 million raw reads (Fig. 1a, Fig. 1d), whereas the output of DRS on the same MinION flow cell is only ~1 million^21,22^. FLEP-seq2 also uses much less input RNA than DRS. DRS requires 500 to 1000 ng polyadenylated RNA^21,22^, while FLEP-seq2 can start with as little as 500 ng total RNA. The full-length information obtained by FLEP-seq2 also enables us to simultaneously track the splicing status, poly(A) site position and poly(A) tail length of each transcript (Fig. 1c). These results demonstrated that FLEP-seq2 is an unbiased, robust, and high-throughput method for measuring poly(A) tail length; therefore, we chose FLEP-seq2 for a comprehensive characterization of poly(A) tail in plants.

**Figure 1.**
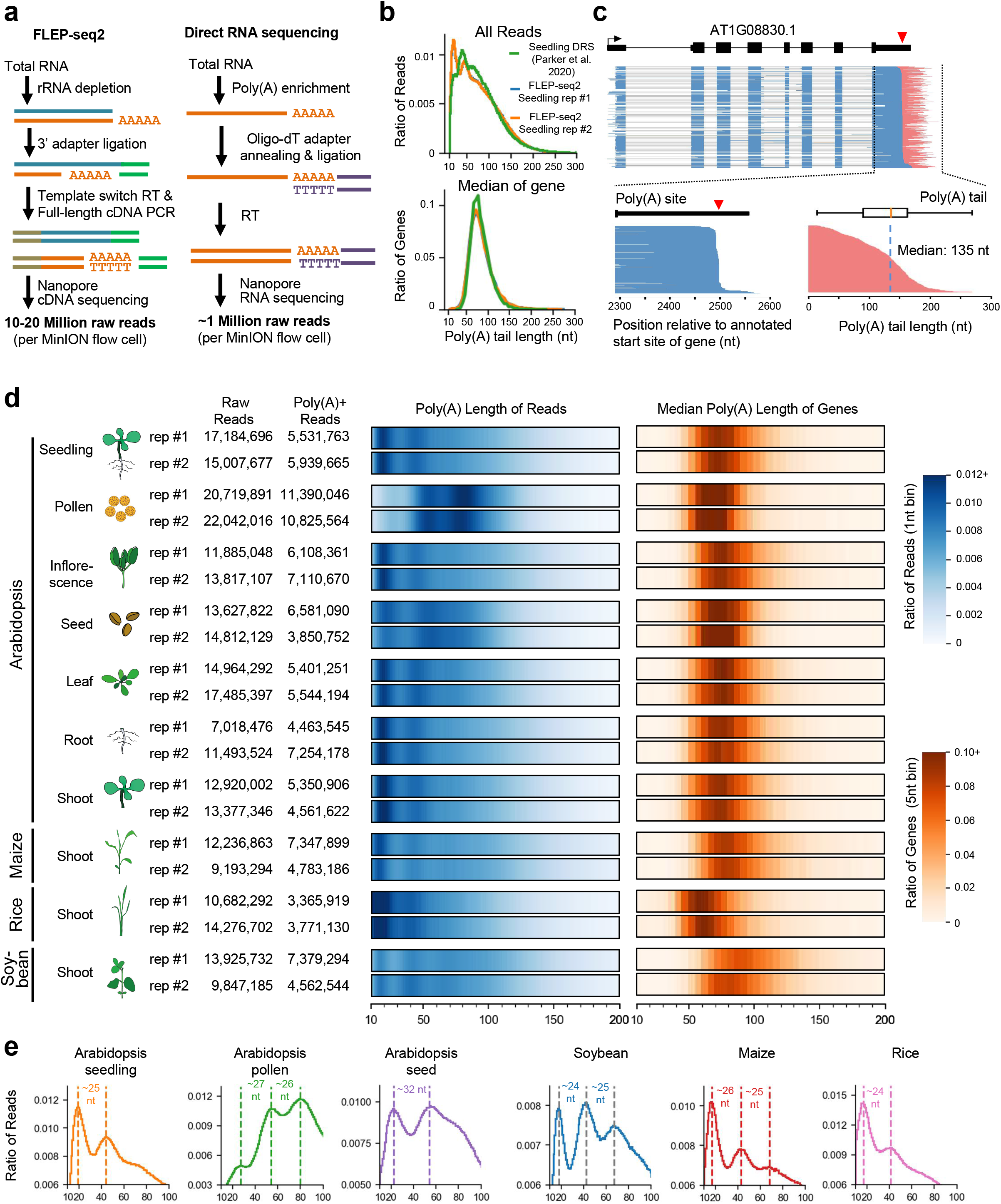
An atlas of plant poly(A)-tail lengths measured by FLEP-seq2. **a**, The schematic diagram of FLEP-seq2 and Direct RNA sequencing (DRS). RT: reverse transcription. **b**, The distribution of global poly(A) tail lengths of transcripts/reads (upper panel, 1 nt bin) and the median poly(A) tail length of genes (bottom panel, 5 nt bin, only genes with at least 20 reads were used) measured by FLEP-seq and DRS. **c**, An example of reads aligned to the AT1G08830 gene in a FLEP-seq2 library (seedling replicate 1). Only polyadenylated reads were shown. **d**, The distribution of global poly(A) tail lengths of transcripts/reads (left panel) and the median poly(A) tail length of genes (right panel) in different tissues and different species. In the analysis of median poly(A) tail length of genes, only genes with at least 20 reads were used. Rep #1: biological replicate 1; rep #2: biological replicate 2. **e**, The peaks of global poly(A) tail length distribution in representative samples.

To establish a comprehensive atlas to investigate the tissue-specificity and evolutionary conservation of poly(A) tail length regulation in plants, we first extracted the total RNA of seven different tissues (seedling, root, shoot, leaf, inflorescence, seed, pollen) of Arabidopsis, as well as the shoot tissues of three important crops: maize, rice, soybean, with two biological replicates for each sample (Fig. 1d). We constructed 20 FLEP-seq2 libraries, each sequenced with a MinION flow cell, which in total yielded ~276 million raw reads and ~121 million poly(A)+ reads (Fig. 1d, Table S1). The median poly(A) length of gene is mainly between 50 nt and 100 nt (Table S2), while the poly(A) length distribution of all reads has two prominent peaks, one at ~20 nt and the other at ~45 nt (Fig. 1 d, Fig. 1e). A peak in ~70 nt can also be observed in some samples, such as soybean and maize (Fig. 1e). These peaks are typically phased with an interval of 25 to 30 nt (Fig. 1e), consistent with the footprint of one PABPC protein^2,6,27–29^, suggesting a large portion of mRNAs are protected by PABPC in plants.

To investigate if the differences in poly(A) tail length among different genes originated from the nascent RNA, we also analyzed the poly(A) tail length of nascent RNA in the nuclei from Arabidopsis, rice, soybean and maize (Table S3). To our surprise, the poly(A) tail lengths of the nuclear RNA are much longer, mainly at 100–200 nt, with the median length of nuclear RNAs are almost twice the size as to those of total RNAs (Fig. S3), and the median poly(A) tail lengths of genes are also considerably longer than those in total RNAs (Fig. 2a). We previously reported that a large portion of plant nascent transcripts are fully transcribed, polyadenylated, yet incompletely spliced, and these intron-containing nascent RNAs are presumably tethered to the chromatin until they complete splicing^5^. Based on this model, the transcripts with introns detected in total RNAs are mostly the incompletely spliced RNAs in the nuclear. Indeed, we observed that the intron-containing transcripts in total RNA have longer poly(A) tails than the fully spliced ones (Fig. 2b, Fig. 2c, Fig. S4a), and the poly(A) tail lengths of these transcripts with introns from total RNA are similar to those from nuclear RNA (Fig. 2b, Fig. 2d, Fig. S4b). Interestingly, in the nuclei, poly(A) tail lengths of the intron-containing transcripts and fully-spliced transcripts are largely the same (Fig. 2b), suggesting shortening of poly(A) tail is a rapid process that occurs after splicing is completed and before the transported mRNA is stablished in the cytoplasm.

**Figure 2.**
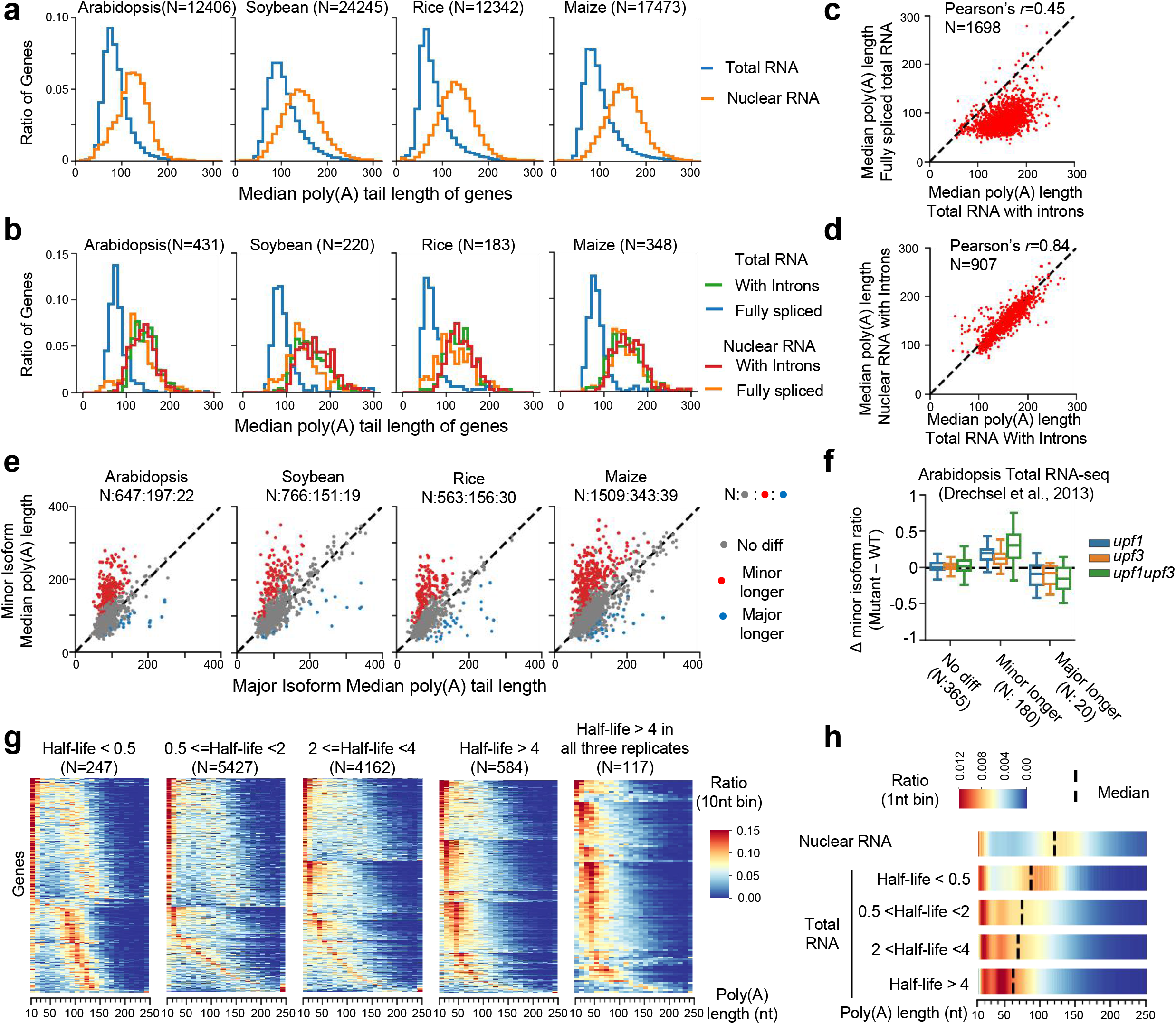
Nuclear Poly(A) tails are longer than tails in cytoplasm. **a**, The distribution of median poly(A) tail lengths of nuclear and total RNA from different plant species (5 nt bin). Only genes with at least 20 detected reads in both nuclear and total RNA libraries were used. **b**, The distribution of median poly(A) tail lengths of fully spliced transcripts and intron-containing transcripts (with introns) from nuclear and total RNA in different plant species (5 nt bin). Only genes with at least 20 fully spliced reads and 20 intron-containing reads in both nuclear and total RNA libraries were used. **c**, The comparison of median poly(A) tail lengths between fully spliced transcripts and intron-containing (with introns) transcripts in Arabidopsis seedling samples. Only the transcripts/reads spanning all annotated introns were used, and only genes with at least 20 fully spliced reads and 20 intron-containing reads are used. **d**, The comparison of median poly(A) tail lengths of intron-containing transcripts between nuclear and total RNA in Arabidopsis seedling. Only the transcripts/reads spanning all annotated introns were used, and only genes with at least 20 intron-containing transcripts in both nuclear and total RNA libraries were used. **e**, The comparison of median poly(A) tail lengths between minor and major alternative-splicing (exclude intron-retention) generating isoforms. Only isoforms with at least 20 reads were used. For each gene, the isoform with the highest expression was designed as major isoforms. All other isoforms were designed as minor isoforms and were compared to the major isoform. The numbers of isoforms showing no differential, longer, and shorter poly(A) tails were separated with “:” and labeled above each figure. **f**, Boxplot showing the distribution of Δ minor isoform ratio (minor/[minor+major]) of mutants. The sum of the number of minor and major isoform reads in each sample is required to be more than 10. **g**, Heatmap plot showing the poly(A) tail length distribution of genes with different half-lives. The mRNA half-life data of seedling was reported in previous paper (Szabo et al., 2020). Each row in the plot represents the poly(A) tail length distribution of one gene. Only genes with at least 50 reads were used. **h**, The bulk poly(A) tail length distribution of genes with different mRNA half-lives in Arabidopsis. N: gene number.

In addition, we found that the poly(A) tail length of about 1/5 of alternatively spliced isoforms (exclude intron retention) are significantly longer than those of the corresponding major isoforms (the most expressed isoforms) in all detected four species (Fig. 2e). Using public Arabidopsis RNA seq data^30^, we found that most of these isoforms, e.g., an alternative 3’ splicing site event generating isoform in AT3G01480 (Fig. S5), are upregulated in *up frameshift 1* (*upf1*) *upf3* mutant (Fig. 2f), which disrupts the cytoplasmic nonsense-mediated decay (NMD) pathway. This suggested that most of them are the targets of the NMD pathway. NMD is a cytoplasmic mRNA surveillance mechanism which primarily recognizes target RNA during the first round of translation and mediates rapid degradation of their targets^31,32^. Based on this model, for these NMD-targeted isoforms, the detected transcripts/reads should mainly be newly synthesized RNAs which are still in the nucleus and haven’t undergone the first round of translation. Consistent with this, these isoforms are enriched in nuclear (Fig. S6), and their poly(A) tail lengths are consistent with those of the transcripts with introns (Fig. S7). All these results indicated that nuclear nascent RNA has a long poly(A) tail and a global deadenylation of mature mRNAs occurs in the cytoplasm of plant cells.

Transcripts from different genes have poly(A) tails of distinct lengths (Fig. 1d, right panel). However, the poly(A) tails of nuclear nascent RNAs are generally different from those of total RNAs (Fig. 2b), indicating that these intergenic length differences of poly(A) tail in steady-state transcripts should be largely determined by the cytoplasmic deadenylation step. Using the reported genome-wide dataset of mRNA half-lives in Arabidopsis^33^, we found that the poly(A) tails of the most unstable transcripts, i.e., the transcripts from genes with shortest mRNA half-lives, are mainly 70–150 nt, similar to or slightly shorter than those of nuclear RNAs (Fig. 2g, 2h). For these genes, a large number of transcripts with short poly(A) tails in 10–20 nt are also detected, while the transcripts with poly(A) tails in 20–70 nt are few, implying that they undergo a rapid deadenylating step that shortens the poly(A) in the cytoplasm. On the contrary, the poly(A) tail lengths of the stable transcripts, especially for the most stable transcripts, are distinctly shorter than those of nuclear nascent RNAs, and usually enriched in the range of 20 to 80 nt with phased peaks presumably due to PABPC-binding (Fig. 2g, 2h). These results suggest stable mRNAs initially undergo cytoplasmic deadenylation but are subsequently protected by PABPs against further deadenylating and decay. These results indicated the different poly(A) tail lengths that we observed for different genes could be partly explained by the regulation of differential deadenylation.

Next, we investigated whether there is tissue-specific regulation of poly(A) tail length. Although the poly(A) tail length distributions are similar among most tissues, the pattern in pollen and seed are distinct (Fig. 1d, Fig. 3a). The poly(A) tails of transcripts in pollen and seed, especially in pollen, are mainly composed of medium size, with few shortest and longest poly(A) tails (Fig. 1d, Fig. 3a), which is reminiscent of the feature of stable transcripts in other tissues (Fig. 2g). The poly(A) tail distribution of pollen RNAs has three peaks with a ~26 nt phase interval (Fig. 1e). Moreover, different from other tissues in which the poly(A) tails are enriched at 20–30 nt and 40– 60 nt, the poly(A) tails of pollen are mainly at 40–60 nt and 70–90 nt (Fig. 1d, Fig. 3a), suggesting that the transcripts in pollen are potentially bound and protected by more PABPs. The poly(A) tails of mRNAs with short half-lives in seedling also enriched at 40–90 nt and has two peaks at 40–60 nt and 70–90 nt in pollen (Fig. S8a, S8b), implying that many transcripts which are rapidly degraded in other tissues are also protected in pollen. Similar to pollen, poly(A) tail distribution in seed also exhibits longer tails, although the pattern is less obvious than in pollen (Fig. 1d, Fig. 1e, Fig. 3a). These results indicated a stronger protection by PABPs in pollen and seed, consistent with the previous reports that the mRNAs in pollen are usually stable^34^ and many mRNAs are stored in pollen and seed for the germination of pollen and seed, respectively ^35,36^.

**Figure 3.**
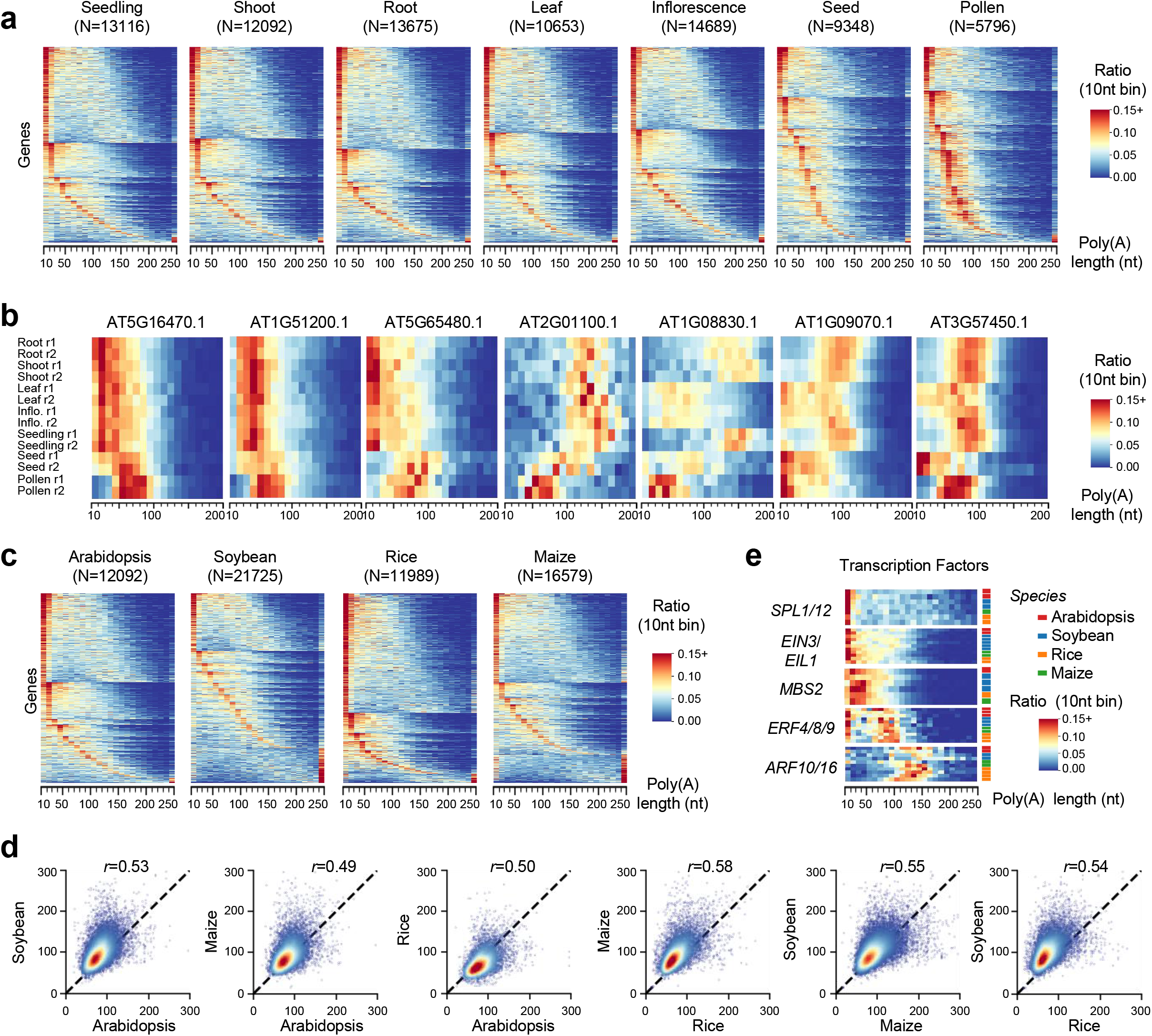
Tissue-specific and evolutionarily-conserved regulation of poly(A) tail length in plants. **a**, Heatmap plot showing the poly(A) tail length distribution of genes in different tissues. Only genes with at least 50 reads were used. **b**, Examples of genes showing differential poly(A) tail length distribution in different tissues. Inflo.: Inflorescence. r1: biological replicate 1; r2: biological replicate 2. **c**, Heatmap plot showing the poly(A) tail length distribution of genes in different species. Only genes with at least 50 reads were used. **d**, The correlation of the median poly(A) tail length of orthologous gene pair in different species. Only genes with more than 50 reads were used. The Pearson’s *r* values were labeled above each figure. **e**, Examples of the poly(A) tail length distributions of homologous genes among different species. N: gene number.

Besides pollen and seed which are distinctly different from other tissues in poly(A) tail length distribution, the correlation coefficients of the median poly(A) tails of genes among the other five tissues are also lower than the correlation coefficients between two biological replicates from the same tissue (Fig. S9a). The heatmap plot of the median poly(A) tail length of genes also showed similar results (Fig. S9b). These results suggested that the poly(A) tails of many genes are tissue-specifically regulated. Consistent with this finding, a large number of genes (range between 250 and 1665) showing significantly differentially regulated poly(A) tails were identified in each pair of tissues, compared to only few (8 to 57) differential genes identified from the random data generated by shuffling the samples of two compared tissues (Fig. S9c). For example, the poly(A) tails of AT5G16470.1, AT1G51200.1, AT5G65480.1 are enriched in 10–50 nt length in most tissues, and the poly(A) tail of AT2G01100.1 are enriched in 100–200 nt length, but all of them are enriched in 50–100 nt length in pollen (Fig. 3b). The poly(A) tail of AT1G08830.1 and AT1G09070.1 are differentially shorter in leaf and inflorescence than those in seedling, shoot and root (Fig. 3b). And the poly(A) tails of AT3G57450.1 are shorter in leaf than those in seedling, shoot, root and inflorescence (Fig. 3b).

The poly(A) length distributions of different species are quite similar, but also show some differences (Fig. 1d, Fig. 3c). Compared to Arabidopsis shoot, rice shoot has more transcripts with an extremely short tail (Fig. 1d), and more genes have a peak at 10–20 nt in poly(A) distribution (Fig. 3c), and thus the median poly(A) tail lengths of genes are shorter (Fig. 1d). In contrast, maize and soybean shoot, especially for soybean shoot, has fewer transcripts with an extremely shorter tail, but has more transcripts with a longer tail (Fig. 1d, Fig. 3c), and showed a higher peak in 60– 80 nt (Fig. 1d, Fig. 1e), which might represent transcripts protected by three PABPs. Despite these differences, the poly(A) tail lengths of homologous genes are significantly correlated among different species (*r*: 0.49 to 0.58) (Fig. 3d). Take transcription factors as examples, the poly(A) tails of *SQUAMOSA PROMOTER BINDING PROTEIN-LIKE 1* (*SPL1*) and *SPL12*, which act redundantly in thermotolerance at the reproductive stage^37^, has a broader distribution in length; *ETHYLENE INSENSITIVE 3* and its closest homolog *EIN3-LIKE1* (*EIL1*), two key regulators in ethylene signal transduction pathway, have more poly(A) tails in 20–100 nt; the poly(A) tails of *METHYLENE BLUE SENSITIVITY 2* (*MBS2*), a mediator of singlet oxygen responses^38^, enriched at 20–50 nt; the poly(A) tails of *ETHYLENE RESPONSE FACTOR 4* (*ERF4*) / *ERF8* / *ERF9* have two peaks in length, one at 50–100 nt and the other at 10–20 nt; and the poly(A) tails of *AUXIN RESPONSE FACTOR 10* (*ARF10*) / *ARF16* are mainly higher than 100–nt (Fig. 3e). These results suggested that the poly(A) tail lengths of orthologous genes are relatively conserved among different plant species, and thus might be selected under evolutionary pressure.

Our data showed that the poly(A) tail lengths can be different among different genes but highly correlated among different tissues for the same gene (Fig. S9a, *r* ⩾ 0.8 for most tissue pairs except for pollen and seed), and are evolutionarily conserved in different species. These results indicate that they are tightly controlled in gene-specific ways in plants, thus may reflect their critical roles in gene regulation. The poly(A) tails of total RNA are significantly shorter than those of nuclear RNA, indicating that the poly(A) lengths of steady-state mRNA are largely dependent on the cytoplasmic deadenylation process. If deadenylation rate is uniform for a given gene, the poly(A) tail length distribution should be broad and flat, such as the pattern of *SPL1/SPL12* genes shown in Fig. 3e. However, the poly(A) tail of many plant transcripts, especially for stable RNAs, have peaks at 20–60 nt, indicating that they first undergo rapid deadenylation and then be protected when poly(A) tails become 20–60 nt. This is consistent with the dual roles of PABPC and the biphasic deadenylation model reported in animals and yeast — PABPC can stimulate the deadenylation of long poly(A) tails via binding to the deadenylase complexes but blocking precocious decay^1,2,27,28^. However, the homologs of the executor which initially trim the long poly(A) tails of nascent RNA in this model, PAN2/PAN3^2,27^, although conserved in animals and yeast, haven’t been identified in the Arabidopsis genome^39^, implying other deadenylase complexes may replace them to perform the initial trimming. Besides, the poly(A) tail of some transcripts, especially for those with short half-lives, are longer and rarely in 20–60 nt, but usually are also enriched in 10–20 nt (Fig 2g, 3a, 3c), a size that may be short enough to loosen or lost their association with PABP and has been reported to prefer for being uridylated and decay^24,28,40^, implying an accelerated deadenylation from long poly(A) tail to extremely short, consistent with the canonical mRNA decay model that the poly(A) tail is first shortened to 10–12 nt before further decay from both 5’-3’ and 3’-5’ direction^2^. These results suggested that there could be multiple modes of deadenylation in plants, which are gene-specifically regulated, highlighting the importance of profiling poly(A) lengths of different genes.

It has been reported that the poly(A) tail lengths are globally changed in specific animal systems, such as the oocyte-to-embryo development stage, a model system to study the function of poly(A) tail^1,2,13,16^. Here, our data highlights that pollen and seed show distinct poly(A) tail length distribution among plant tissues. The poly(A) tails of pollen and seed RNAs are enriched in 40– 90 nt, thus might be protected by more PABPC proteins and serve the purpose of storing mRNA in these two tissues^34–36^. Arabidopsis genome contains eight PABPC genes^8,41^. Previous reports^8,41^ and the public RNA-seq dataset^42^ reveal that *PAB2* and its two closest homologs, *PAB4* and *PAB8*, are highly expressed in a wide range of tissues, while *PAB3, PAB5, PAB6* and *PAB7* are expressed in pollen (Fig. S10). Further study on these pollen-specifically expressed PABs will help explore the unique mechanism of poly(A) length regulation in pollen and its roles in stabilizing mRNA. Finally, the comprehensive landscape of poly(A) tails from various tissues and species will provide an important resource to explore the dynamic regulation of poly(A) length and its roles in controlling gene expression in plants.

## Methods

### Plant materials and RNA isolation

Arabidopsis ecotype Col-0, soybean cultivar Wm82, rice cultivar Nipponbare and maize cultivar B73 are used in this study. For Arabidopsis, plants were grown at 22°C with 16 h of light per 24 hours. Arabidopsis seedlings, shoots and roots were harvested after growing on 1/2 MS plates for 12 days. Arabidopsis leaves were harvested after growing in soil for 30 days, and Arabidopsis inflorescences were harvested after flowering. For rice, soybean and maize, plants were grown at 28°C with 16 hours of light per 24 hours, and 14-day-old shoots were harvested. The nuclear fractions were separated as described^4,5^, and the total RNA and nuclear RNA were extracted using RNAprep Pure Plant Plus Kit (Polysaccharides & Polyphenolics-rich, TIANGEN, DP441) according to the manufacturer’s instructions.

### Library preparation and Nanopore sequencing

FLEP-seq2 libraries were prepared from 500–3000 ng input RNA as previously described^4,5^ with some improvement. In brief, the ribosomal RNA (rRNA) was removed using pan-plant riboPOOLs probes (siTOOLs Biotech) and Dynabeads Myone Streptavidin C1 (Thermo Fisher, #65001) according to the manufacturer’s instruction (siTOOLs Biotech, two-step depletion method). rRNA-depleted RNA was purified by RNA Clean & Concentrator-5 kit (ZYMO, R1013) and then ligated to 50 pmol 3’ adapter (5’-rAppCTGTAGGCACCATCAAT - NH_2_-3’) in a 20 μl reaction containing 1X T4 RNA Ligase Reaction Buffer, 25% PEG8000, 40U Murine RNase Inhibitor (Vazyme, R301-03) and 20U T4 RNA Ligase 2, truncated K227Q (NEB, M0242) for 10 h at 16°C. The product was cleaned up using RNA Clean & Concentrator-5 kit (ZYMO, R1013) and added into 20 μl of reverse transcription and strand-switching reaction containing 100 nM custom primer (5’ - phos/ ACTTGCCTGTCGCTCTATCTTCATTGATGGTGCCTACAG - 3’), 500 μM dNTPs, 1X RT Buffer, 40U Murine RNase Inhibitor (Vazyme, R301-03), 1 μM Strand-Switching Primer (Nanopore, SQK-PCS109) and 200U Maxima H Minus Reverse Transcriptase (Thermo Fisher, EP0752), and then incubated at 90 min 42°C; 10 cycles of (2 min at 50°C; 2 min at 42°C); and 5 min at 85°C, in a thermal cycler. cDNA libraries were amplified with PrimeSTAR GXL DNA polymerase (TaKaRa, R050A) for 10-16 cycles. To minimize PCR bias, PCR cycle number optimization was performed as previously described^4,5^. After PCR, 1 μl of Exonuclease I (NEB, M0293) was added to the reaction mixture and incubated the reaction at 37°C for 15 min, followed by 80°C for 15 minutes. The products were cleaned up with 0.8X Ampure XP beads (Beckman, A63880). For the Nanopore sequencing, 100 fmol amplified cDNA was loaded onto an R9.4 Flow Cell (Oxford Nanopore Technologies) using Sequence-specific cDNA-PCR Sequencing kit (Nanopore, SQK-PCS109) and sequenced on a MinION device for ~48 hours.

### Nanopore data processing

The Nanopore data pre-processing was performed using FLEP-seq pipeline (https://github.com/ZhaiLab-SUSTech/FLEPSeq) as previously described with minor modification about adapter sequence^4,5^. Briefly, the raw Nanopore signals were converted to base sequences by Guppy (v4.0.11, download from Oxford Nanopore Community) with the default parameters (--c dna_r9.4.1_450bps_hac.cfg). The reads with a mean quality score of more than 7 were mapped to genome sequence using Minimap2 (v.2.17-r943-dirty)^43^ with the following parameters: -ax splice --secondary=no -G 12000. The used genome and annotation versions of Arabidopsis, soybean, rice, maize were ARAPROT11 (https://www.arabidopsis.org/), Wm82.gnm4.ann1.T8TQ (https://soybase.org/), MSU7 (http://rice.uga.edu), B73 RefGen_v5 (https://maizegdb.org), respectively. The reads mapped to rDNA, mitochondria and chloroplast genomes were removed by filter_rRNA_bam.py in FLEP-seq pipeline. The 3’ adapters were identified by adapterFinder.py with parameter: --mode 1. The 5’ and 3’ adapter sequences of FLEP-seq2 are different from those of FLEP-seq, and are TTTCTGTTGGTGCTGATATTGCT and ACTTGCCTGTCGCTCTATCTTCATTGATGGTGCCTACAG, respectively. This modification has been integrated into the new version of adapterFinder.py (set ‘--mode 1’ parameter for FLEP-seq2, and ‘--mode 0’ or default for FLEP-seq) in FLEP-seq pipeline. Then, poly(A) tail identification and length estimation were performed by PolyACaller.py and the splicing status of intron was extracted by extract_read_info.py. The transcripts/reads with a predicated poly(A) tail score equal to or more than 10 were identified as polyadenylated transcripts and used for further analysis. Only the reads spanning all introns were used for the analysis of the fully-spliced and intron-containing transcripts, and the intron with a mapping ratio of at least 80% in a read was identified as unspliced intron. To identify orthologs among different species, the protein sequences of all four species were exported to OrthoFinder software^44^ with default parameters.

### Identification of alternative splicing isoform from Nanopore data

We first extracted splicing junctions from reads mapped to a given gene to identify high credible splicing events. Due to the higher base error ratio, the alignment quality of nanopore reads near the splice site is relatively poor, and the end positions of splicing junctions might be wrong. Thus, if both ends of multiple detected splicing junctions are close (within 10 nt), they might be generated from wrong alignments, and only the splicing junction with the most supported reads was remained. The set of overlapping splicing junctions represents a group of alternative splicing (AS) events. For a given splicing junction (*J*), the percent spliced in (PSI) value of *J* equal to *n1*/[*n1*+*n2*]; *n1*: the number of reads specifically supported *J*; *n2*: the number of reads specifically supported the splicing junctions overlapping with *J*. The splicing junction with less than 20 supported reads or with a PSI value lower than 2% were removed. Multiple introns of one gene might undergo AS, and thus one gene could contain multiple AS groups. Therefore, we second joined these AS junctions to AS isoforms based on the supporting reads. For a given gene, the reads spanning all AS groups were extracted. The splicing statuses of all AS junctions in each read were extracted, and the intron-containing transcripts/reads were removed. Each remained read was explicitly derived from one kind of isoform, and all supported AS junction combinations/isoforms as well as the number of supported reads were calculated. The isoforms with more than 20 supported reads were identified as highly creditable isoforms, and the poly(A) tails of the reads specifically supporting them were used for the poly(A) tail length analysis of isoforms

### Isoform quantification

For a given gene, unique representative splicing junction of each isoform was extracted. Considering that the sequencing depth of the 3’ end of a gene is usually higher, if one isoform contains two or more unique representative splicing junctions, only the most downstream one was used. For each sample, the number of reads supported the unique representative splicing junction of each isoform (*n_j_*) was extracted, and the minor/major isoform ratio was calculated by *n_j_minor_/n_j_major_*.

### Identification of genes showing differential poly(A) tail length

For comparison of two groups (e.g., tissue S and tissue R, or major isoforms S and minor isoforms R) with biological replicates (e.g., S1, S2, R1 and R2), for each gene, the poly(A) tail lengths of gene-derived reads detected in each sample (*X_S1_, X_S2_, X_R1_, X*_R2_) were extracted, and the medians of poly(A) tail lengths of each sample (M_S1_, M_S2_, M_R1_, M_R2_) were calculated. The difference factor of median (*dm*) was calculated by *dm* = max(min(0, *dm1*), min(0, *dm2*)); *dm1* = min(M_S1_, M_S2_) – max(M_R1_, M_R2_); dm2= min(M_R1_, M_R2_) – max(M_S1_, M_S2_). *X_S1_* and *X_S2_* were merged to *X_S_*, and *X_R1_* and *X_R2_* were merged to *X_R_*. Then, for each gene, a two-sided Mann-Whitney U test between *X_S_* and *X_R_* was performed, and the p-values of all genes were adjusted by Benjamini/Hochberg FDR (False Discovery Rate) method. The genes with an adjusted p-value less than 0.05 and a value *dm* more than 20 were identified as the genes showing differential poly(A) tail length between samples.

### Data Availability

The raw sequencing data generated in this study were deposited in China National Center for Bioinformation with accession PRJCA007575 and in NCBI with accession PRJNA788163 (https://dataview.ncbi.nlm.nih.gov/object/PRJNA788163?reviewer=hlvulfu2lo8062gc5fhuuiupf6 for reviewer link). The poly(A) tail lengths of reads from each gene in each library were recorded in China National Center for Bioinformation with accession OMIX881. The median poly(A) lengths of genes in each library were recorded in Table S2.

**Correspondence and requests for materials** should be addressed to Jixian Zhai.

## Acknowledgments

The group of J.Z. is supported by the National Key R&D Program of China Grant (2019YFA0903903); the Program for Guangdong Introducing Innovative and Entrepreneurial Teams (2016ZT06S172); the Shenzhen Sci-Tech Fund (KYTDPT20181011104005); the Key Laboratory of Molecular Design for Plant Cell Factory of Guangdong Higher Education Institutes (2019KSYS006); and the Stable Support Plan Program of Shenzhen Natural Science Fund Grant (20200925153345004). J.J. is supported by the National Natural Science Foundation of China (32100444); and the Shenzhen Fundamental Research Program (20210317103146009).

## Author Contributions

J.Z., J.J. and W.L. designed the experiments. W.L., J.J., B.L., X.J., Y.S. and Y.L. performed the experiments. J.J., W.L. and Y.Y. analyzed the data. J.Z. oversaw the study. J.J., W.L. and J.Z. wrote the manuscript, and all authors revised the manuscript.

---

**Supplemental Figure 1.**
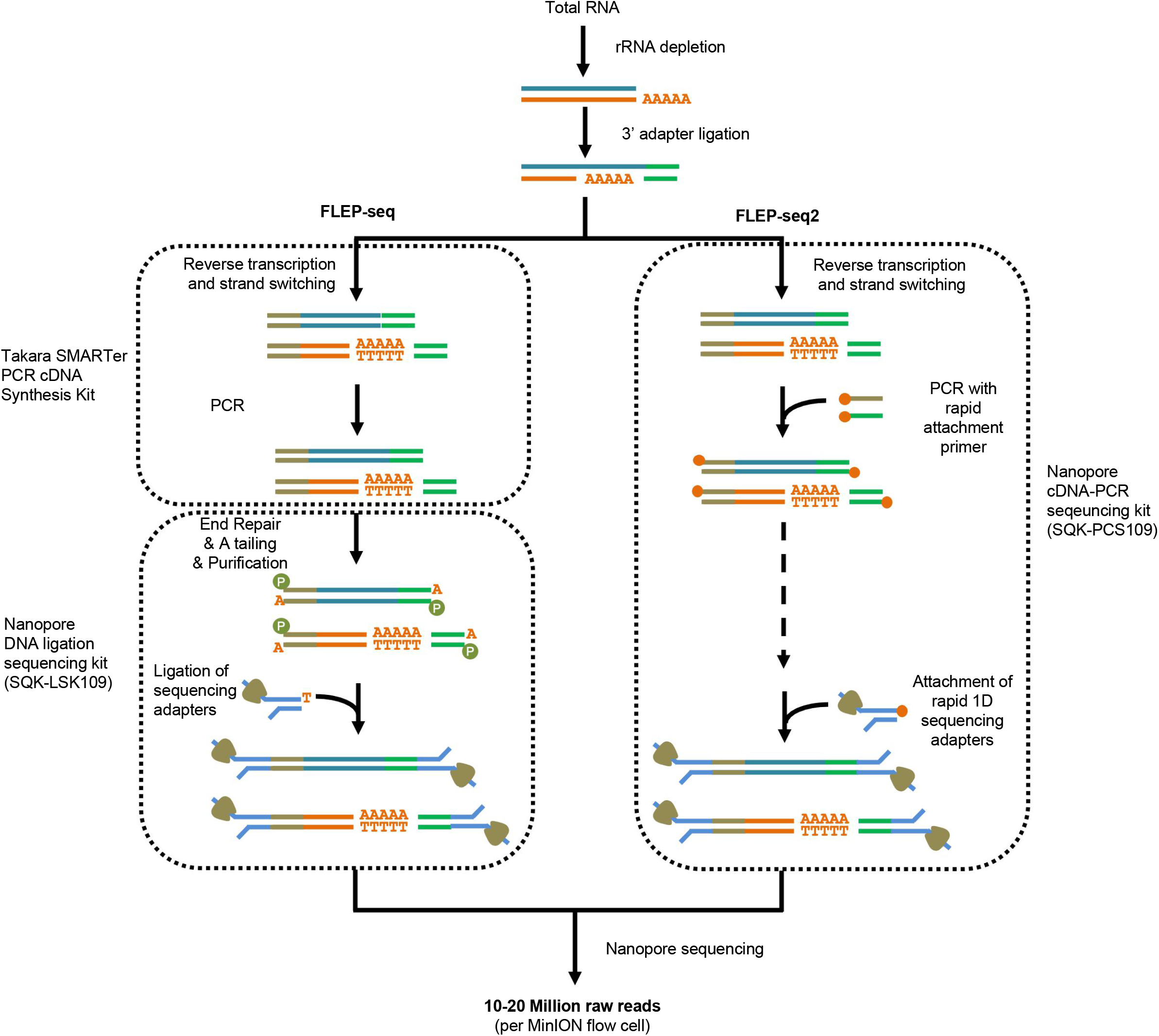
The schematic diagram of FLEP-seq and FLEP-seq2.

**Supplemental Figure 2.**
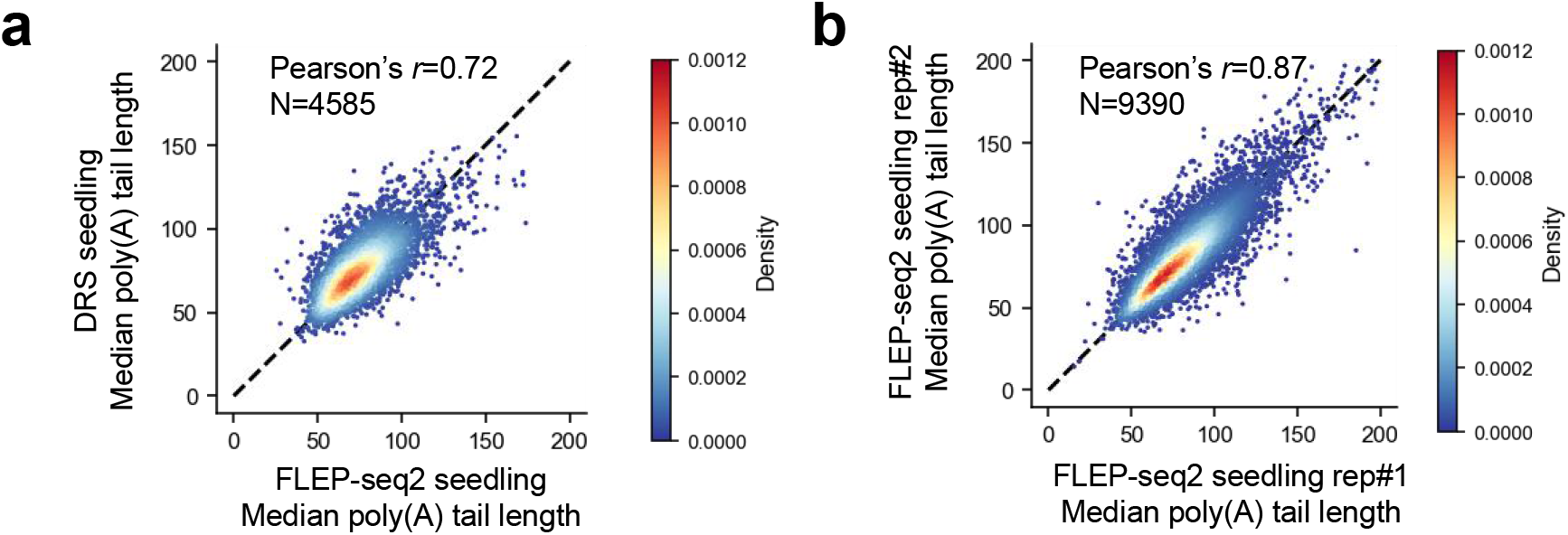
The measured median poly(A) lengths of genes are consistent between FLEP-seq2 and DRS dataset and between different biological replicates of FLEP-seq2. **a**, The correlation of the median poly(A) tail lengths measured by FLEP-seq2 and Direct RNA sequencing (DRS). Only genes with at least 50 reads in both data set were used. **b**, The correlation of the median poly(A) tail lengths measured by FLEP-seq2 in two different biological replicates. Only genes with at least 50 reads in both samples were used.

**Supplemental Figure 3.**
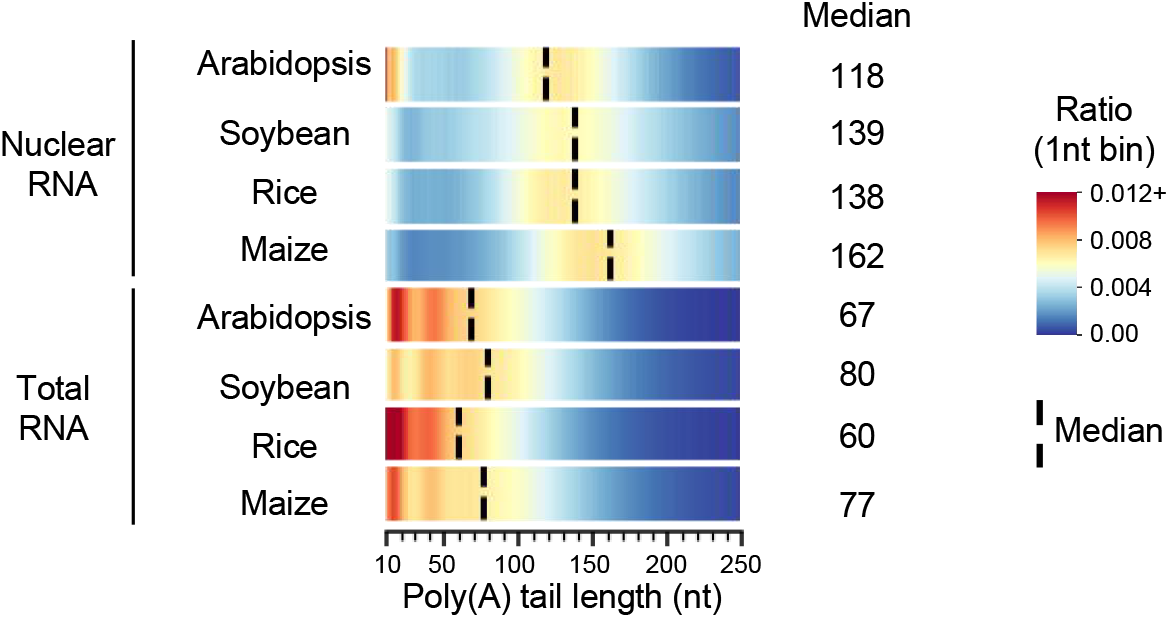
The poly(A) tail length distribution of total RNA and nuclear RNA in different species.

**Supplemental Figure 4.**
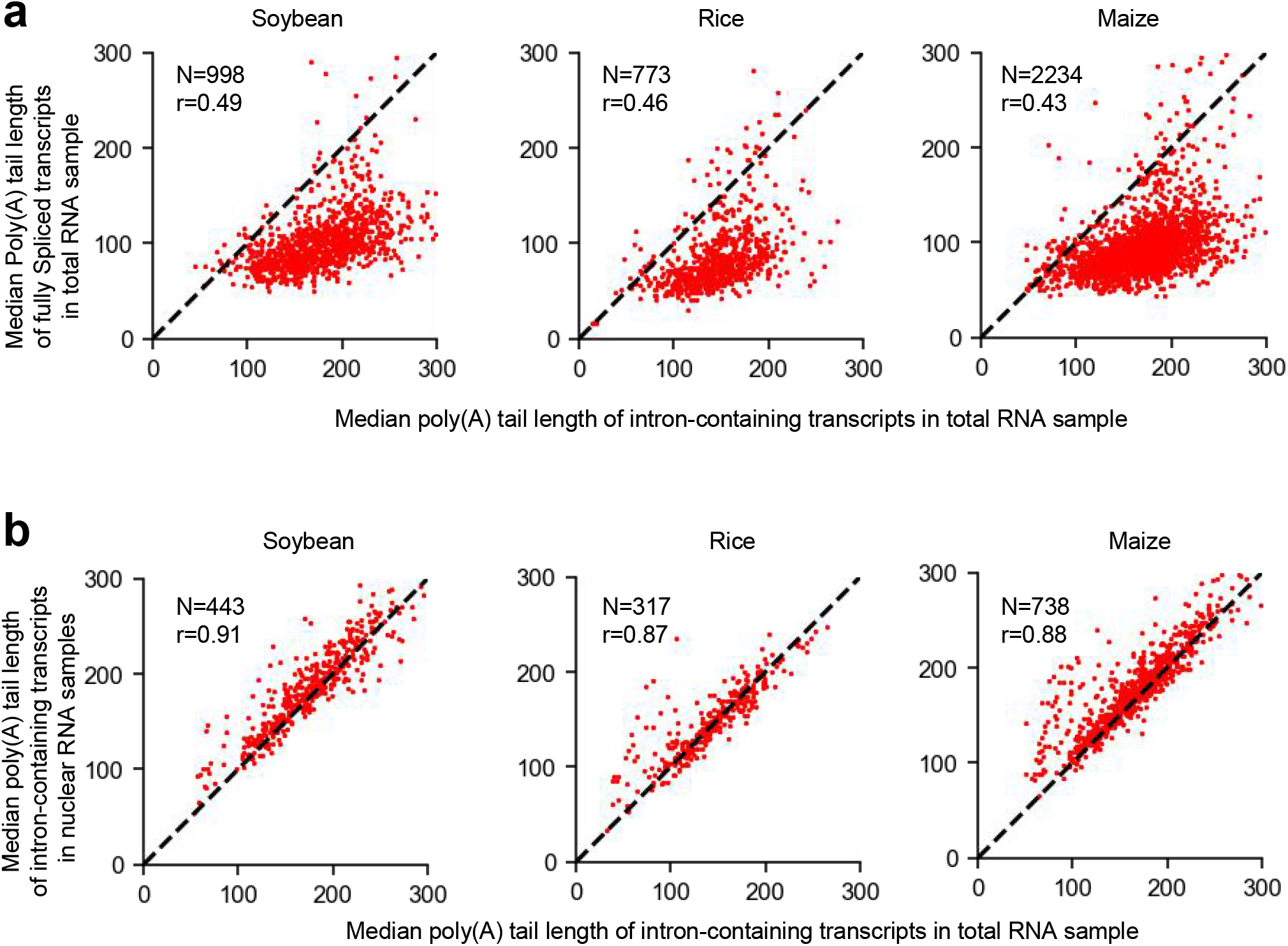
Comparison of the poly(A) tail length of intron-containing transcripts and fully-spliced transcripts. **a**, The comparison of median poly(A) tail lengths between fully spliced transcripts and intron-containing (with introns) transcripts in total RNA samples. Only transcripts spanning all annotated introns were used, and only genes with at least 20 fully spliced reads and 20 intron-containing reads were used. **b**, The comparison of median poly(A) tail lengths of intron-containing transcripts between nuclear and total RNA samples in different species. Only the transcripts spanning all annotated introns were used, and only genes with at least 20 intron-containing transcripts in both nuclear and total RNA libraries were used. N: gene number. The Pearson’s r values were labeled upper each figure.

**Supplemental Figure 5.**
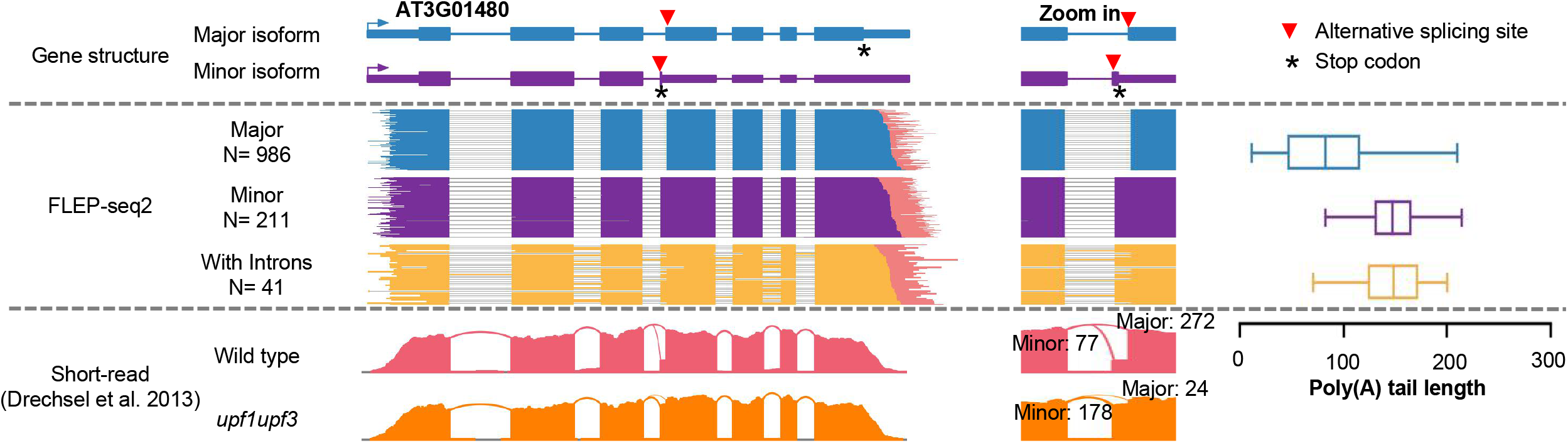
Example of alternative splicing isoforms showing differential poly(A) tail lengths.

**Supplemental Figure 6.**
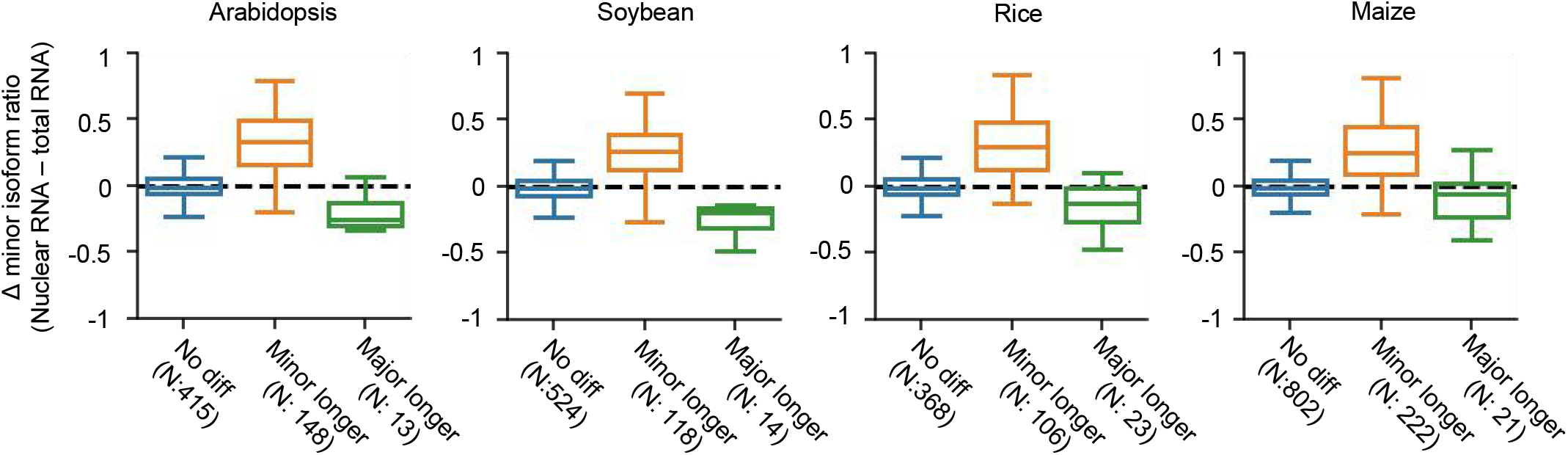
Boxplot showing the distribution of Δ minor isoform ratio (minor/[minor+major]) between nuclear RNAs and total RNAs. The sums of the number of minor and major isoform reads in both nuclear and total RNA samples are required to be more than 10.

**Supplemental Figure 7.**
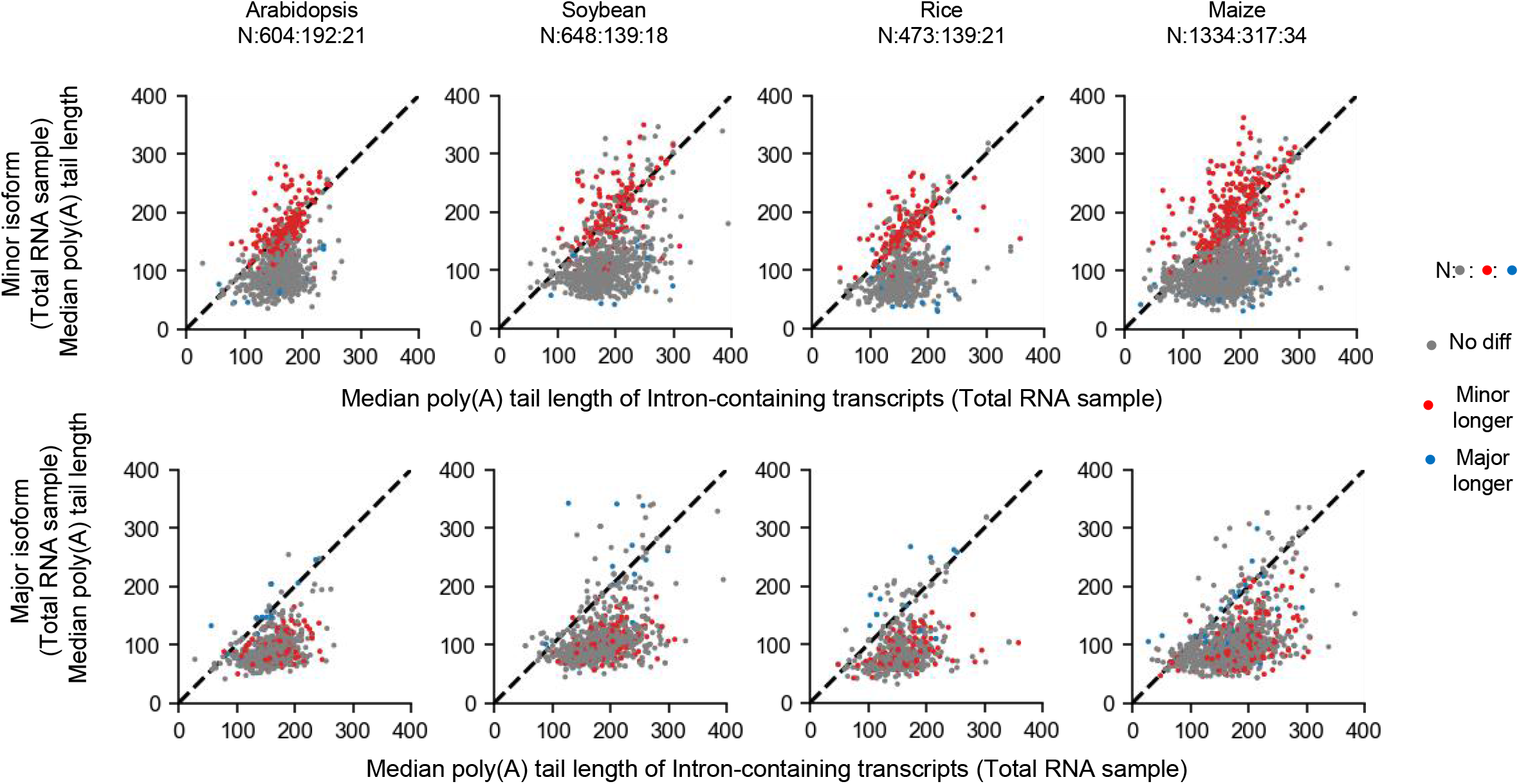
The poly(A) tail lengths of minor isoforms with longer tail than major isoforms are highly consistent with the poly(A) tail of intron-containing transcripts. The comparison of median poly(A) tail lengths between intron-containing transcripts with minor isoforms (upper) or major isoforms (below) in total RNA samples of different species. Only genes with at least 20 minor isoform reads, 20 major isoform reads and 20 intron-containing reads were used. The numbers of isoforms showing no differential, longer, and shorter poly(A) tails were separated with “:” and labeled above each figure.

**Supplemental Figure 8.**
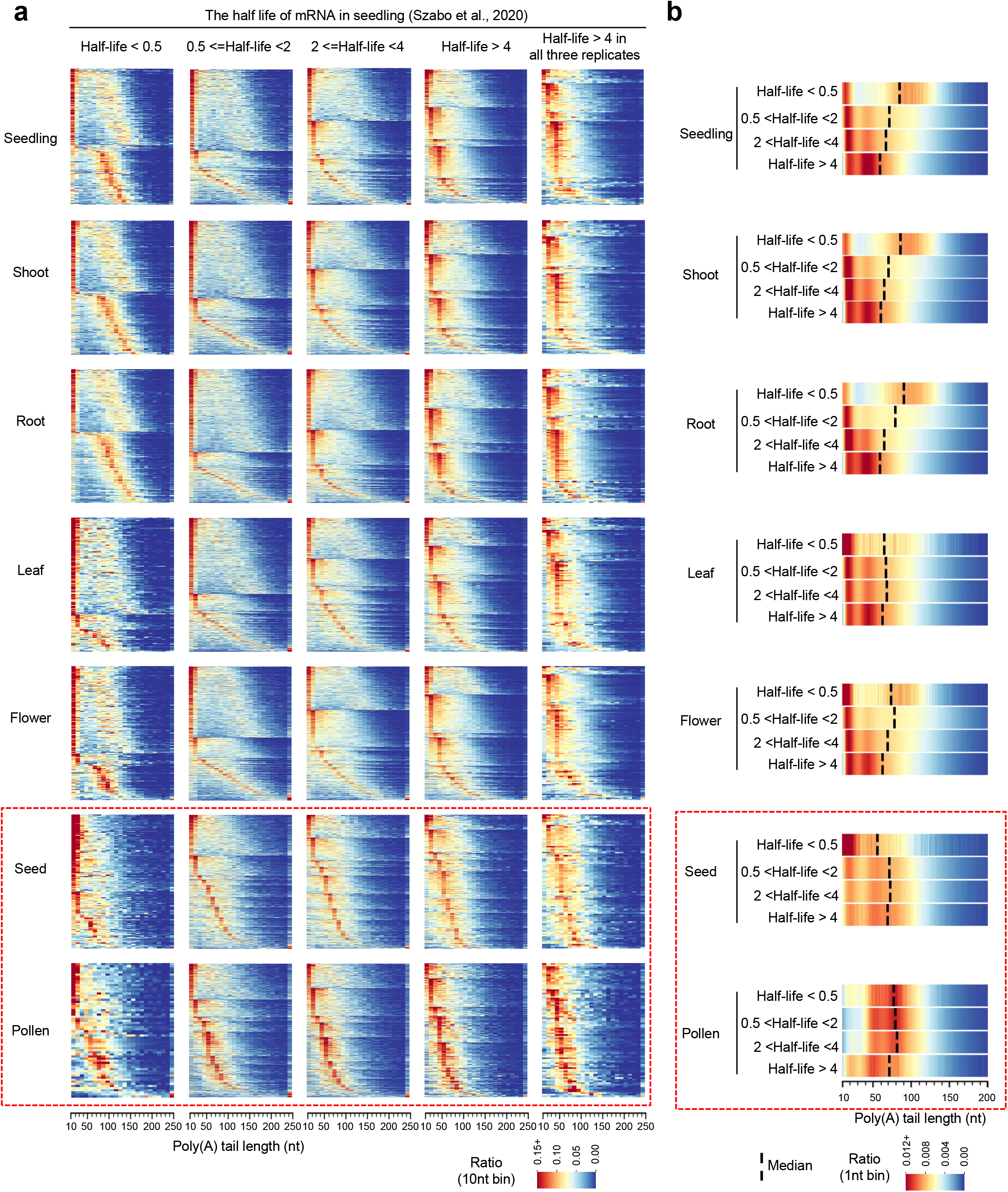
The poly(A) tail lengths of transcripts with short half-lives in seedling are also enriched at 50-100 nt in pollen. **a**, Heatmap plot showing the poly(A) tail length distribution of genes showing different half-lives. The mRNA half-life data of seedling was reported in previous paper (Szabo et al., 2020). Each row in the plot represent the poly(A) tail length distribution a one gene. Only genes with at least 50 reads were used. N: gene number. **b**, The bulk poly(A) tail length distribution of genes with different mRNA half-lives.

**Supplemental Figure 9.**
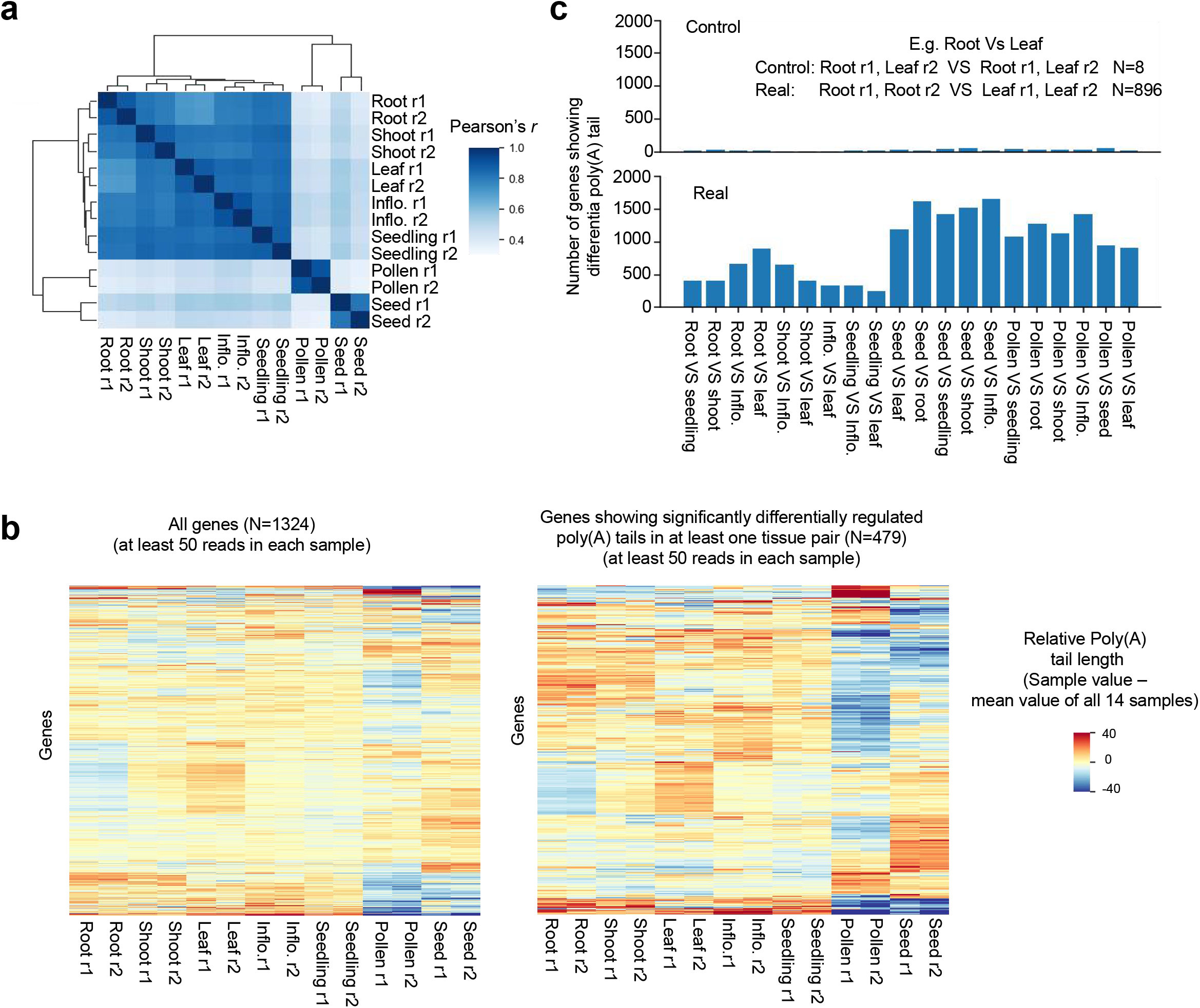
Tissue-specific and evolutionarily-conserved regulation of poly(A) tail length in Arabidopsis. **a**, The median poly(A) tail length correlation matrix of different samples. **b**, Heatmap plot showing the relative median poly(A) tail length of genes among different tissues. **c**, The number of genes identified to show differential poly(A) tail length distribution in different tissues (Real, bottom panel) and in random data (Control, upper panel). Inflo.: Inflorescence. r1: biological replicate 1; r2: biological replicate 2.

**Supplemental Figure 10.**
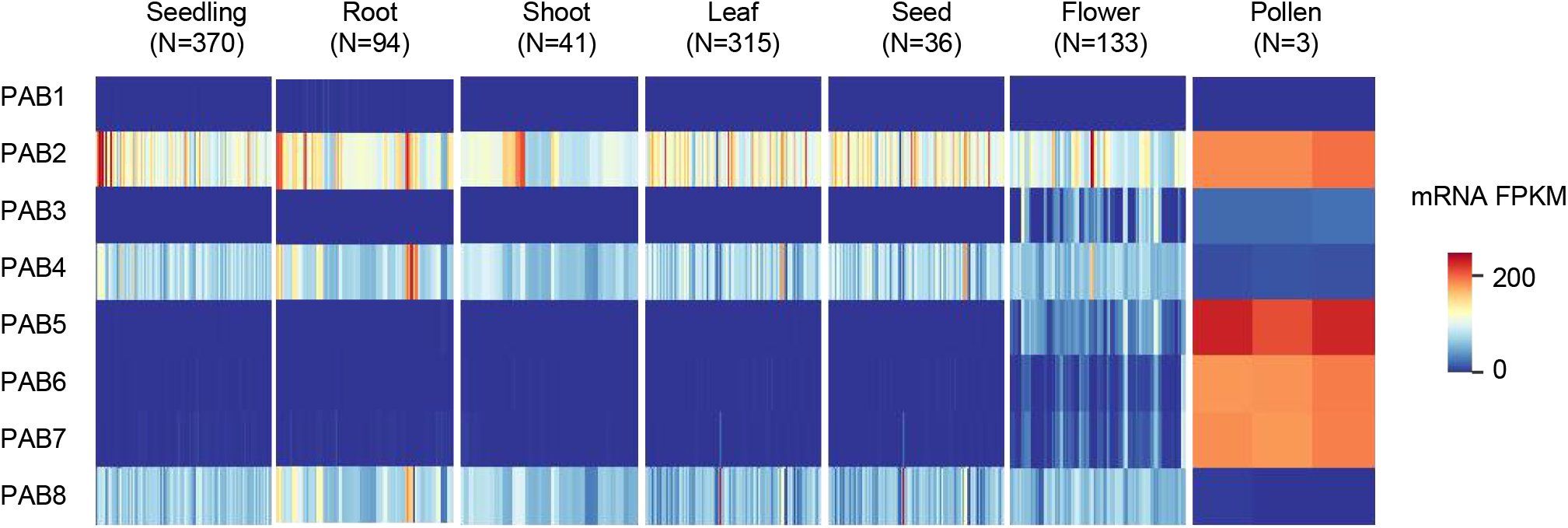
Tissue-specific expression of Arabidopsis genes coding Poly(A) binding proteins (PAB). The gene expression values (Fragments per kilobase per million mapped fragments, FPKM value) of *PABs* in public RNA-seq data are downloaded from http://ipf.sustech.edu.cn/pub/athrna/. N: the number of RNA-seq libraries.

## References

1 Nicholson, A. L. & Pasquinelli, A. E. Tales of Detailed Poly(A) Tails. Trends Cell Biol. 29, 191–200, doi:10.1016/j.tcb.2018.11.002 (2019).

2 Passmore, L. A. & Coller, J. Roles of mRNA poly(A) tails in regulation of eukaryotic gene expression. Nat. Rev. Mol. Cell Biol., doi:10.1038/s41580-021-00417-y (2021).

3 Eckmann, C. R., Rammelt, C. & Wahle, E. Control of poly(A) tail length. Wiley Interdiscip. Rev. RNA 2, 348–361, doi:10.1002/wrna.56 (2011).

4 Long, Y., Jia, J., Mo, W., Jin, X. & Zhai, J. FLEP-seq: simultaneous detection of RNA polymerase II position, splicing status, polyadenylation site and poly(A) tail length at genome-wide scale by single-molecule nascent RNA sequencing Nat. Protoc. 16, 4355–4381, doi:10.1038/s41596-021-00581-7 (2021).

5 Jia, J. et al. Post-transcriptional splicing of nascent RNA contributes to widespread intron retention in plants. Nat. Plants 6, 780–788 doi:10.1038/s41477-020-0688-1 (2020).

6 Lima, S. A. et al. Short poly(A) tails are a conserved feature of highly expressed genes. Nat. Struct. Mol. Biol. 24, 1057–1063, doi:10.1038/nsmb.3499 (2017).

7 Kühn, U. & Wahle, E. Structure and function of poly(A) binding proteins. Biochim. Biophys. Acta. 1678, 67–84, doi:10.1016/j.bbaexp.2004.03.008 (2004).

8 Zhao, T. et al. Impact of poly(A)-tail G-content on Arabidopsis PAB binding and their role in enhancing translational efficiency. Genome Biol. 20, 189, doi:10.1186/s13059-019-1799-8 (2019).

9 Goldstrohm, A. C. & Wickens, M. Multifunctional deadenylase complexes diversify mRNA control. Nat. Rev. Mol. Cell Biol. 9, 337–344, doi:10.1038/nrm2370 (2008).

10 Eisen, T. J. et al. The Dynamics of Cytoplasmic mRNA Metabolism. Mol. Cell 77, 786–799 e710, doi:10.1016/j.molcel.2019.12.005 (2020).

11 Vi, S. L. et al. Target specificity among canonical nuclear poly(A) polymerases in plants modulates organ growth and pathogen response. Proc. Natl. Acad. Sci. U S A 110, 13994–13999, doi:10.1073/pnas.1303967110 (2013).

12 Hunt, A. G. mRNA 3’ end formation in plants: Novel connections to growth, development and environmental responses. Wiley Interdiscip. Rev. RNA 11, e1575, doi:10.1002/wrna.1575 (2020).

13 Subtelny, A. O., Eichhorn, S. W., Chen, G. R., Sive, H. & Bartel, D. P. Poly(A)-tail profiling reveals an embryonic switch in translational control. Nature 508, 66–71 doi:10.1038/nature13007 (2014).

14 Udagawa, T. et al. Bidirectional control of mRNA translation and synaptic plasticity by the cytoplasmic polyadenylation complex. Mol. Cell 47, 253–266, doi:10.1016/j.molcel.2012.05.016 (2012).

15 Chang, H., Lim, J., Ha, M. & Kim, V. N. TAIL-seq: Genome-wide Determination of Poly(A) Tail Length and 3’ End Modifications. Mol. Cell 53, 1044–1052 doi:10.1016/j.molcel.2014.02.007 (2014).

16 Lim, J., Lee, M., Son, A., Chang, H. & Kim, V. N. mTAIL-seq reveals dynamic poly(A) tail regulation in oocyte-to-embryo development. Genes Dev. 30, 1671–1682 doi:10.1101/gad.284802.116 (2016).

17 Harrison, P. F. et al. PAT-seq: a method to study the integration of 3’-UTR dynamics with gene expression in the eukaryotic transcriptome. RNA 21, 1502–1510, doi:10.1261/rna.048355.114 (2015).

18 Woo, Y. M. et al. TED-Seq Identifies the Dynamics of Poly(A) Length during ER Stress. Cell Rep. 24, 3630–3641, doi:10.1016/j.celrep.2018.08.084 (2018).

19 Legnini, I., Alles, J., Karaiskos, N., Ayoub, S. & Rajewsky, N. FLAM-seq: full-length mRNA sequencing reveals principles of poly(A) tail length control. Nat. Methods 16, 879–886 doi:10.1038/s41592-019-0503-y (2019).

20 Liu, Y., Nie, H., Liu, H. & Lu, F. Poly(A) inclusive RNA isoform sequencing (PAIso-seq) reveals wide-spread non-adenosine residues within RNA poly(A) tails. Nat. Commun. 10, 5292 doi:10.1038/s41467-019-13228-9 (2019).

21 Parker, M. T. et al. Nanopore direct RNA sequencing maps the complexity of Arabidopsis mRNA processing and m6A modification. eLife 9, e49658 doi:10.7554/eLife.49658 (2020).

22 Workman, R. E. et al. Nanopore native RNA sequencing of a human poly(A) transcriptome. Nat. Methods 16, 1297–1305, doi:10.1038/s41592-019-0617-2 (2019).

23 Parker, M. T. et al. Widespread premature transcription termination of Arabidopsis thaliana NLR genes by the spen protein FPA. Elife 10, doi:10.7554/eLife.65537 (2021).

24 Scheer, H. et al. The TUTase URT1 connects decapping activators and prevents the accumulation of excessively deadenylated mRNAs to avoid siRNA biogenesis. Nat. Commun. 12, 1298, doi:10.1038/s41467-021-21382-2 (2021).

25 Zuber, H. et al. Uridylation and PABP Cooperate to Repair mRNA Deadenylated Ends in Arabidopsis. Cell Rep. 14, 2707–2717, doi:10.1016/j.celrep.2016.02.060 (2016).

26 Wu, X., Wang, J., Wu, X., Hong, Y. & Li, Q. Q. Heat Shock Responsive Gene Expression Modulated by mRNA Poly(A) Tail Length. Front. Plant Sci. 11, doi:10.3389/fpls.2020.01255 (2020).

27 Schafer, I. B. et al. Molecular Basis for poly(A) RNP Architecture and Recognition by the Pan2-Pan3 Deadenylase. Cell 177, 1619–1631, doi:10.1016/j.cell.2019.04.013 (2019).

28 Yi, H. et al. PABP Cooperates with the CCR4-NOT Complex to Promote mRNA Deadenylation and Block Precocious Decay. Mol. Cell 70, 1081–1088 doi:10.1016/j.molcel.2018.05.009 (2018).

29 Webster, M. W. et al. mRNA Deadenylation Is Coupled to Translation Rates by the Differential Activities of Ccr4-Not Nucleases. Mol. Cell 70, 1089–1100 doi:10.1016/j.molcel.2018.05.033 (2018).

30 Drechsel, G. et al. Nonsense-mediated decay of alternative precursor mRNA splicing variants is a major determinant of the Arabidopsis steady state transcriptome. Plant Cell 25, 3726–3742, doi:10.1105/tpc.113.115485 (2013).

31 Kervestin, S. & Jacobson, A. NMD: a multifaceted response to premature translational termination. Nat. Rev. Mol. Cell Biol. 13, 700–712, doi:10.1038/nrm3454 (2012).

32 Kurosaki, T., Popp, M. W. & Maquat, L. E. Quality and quantity control of gene expression by nonsense-mediated mRNA decay. Nat. Rev. Mol. Cell Biol. 20, 406–420, doi:10.1038/s41580-019-0126-2 (2019).

33 Szabo, E. X. et al. Metabolic Labeling of RNAs Uncovers Hidden Features and Dynamics of the Arabidopsis Transcriptome. Plant Cell 32, 871–887 doi:10.1105/tpc.19.00214 (2020).

34 Ylstra, B. & McCormick, S. Analysis of mRNA stabilities during pollen development and in BY2 cells. Plant J. 20, 101–108, doi:10.1046/j.1365-313x.1999.00580.x (1999).

35 Bai, B. et al. Seed-Stored mRNAs that Are Specifically Associated to Monosomes Are Translationally Regulated during Germination. Plant Physiol. 182, 378–392, doi:10.1104/pp.19.00644 (2020).

36 Hao, H., Li, Y., Hu, Y. & Lin, J. Inhibition of RNA and protein synthesis in pollen tube development of Pinus bungeana by actinomycin D and cycloheximide. New Phytol. 165, 721–729, doi:10.1111/j.1469-8137.2004.01290.x (2005).

37 Chao, L. M. et al. Arabidopsis Transcription Factors SPL1 and SPL12 Confer Plant Thermotolerance at Reproductive Stage. Mol. Plant 10, 735–748, doi:10.1016/j.molp.2017.03.010 (2017).

38 Shao, N., Duan, G. Y. & Bock, R. A mediator of singlet oxygen responses in Chlamydomonas reinhardtii and Arabidopsis identified by a luciferase-based genetic screen in algal cells. Plant Cell 25, 4209–4226, doi:10.1105/tpc.113.117390 (2013).

39 Chantarachot, T. & Bailey-Serres, J. Polysomes, Stress Granules, and Processing Bodies: A Dynamic Triumvirate Controlling Cytoplasmic mRNA Fate and Function. Plant Physiol. 176, 254–269, doi:10.1104/pp.17.01468 (2018).

40 Yu, S. & Kim, V. N. A tale of non-canonical tails: gene regulation by post-transcriptional RNA tailing. Nat. Rev. Mol. Cell Biol. 21, 542–556, doi:10.1038/s41580-020-0246-8 (2020).

41 Belostotsky, D. A. Unexpected Complexity of Poly(A)-Binding Protein Gene Families in Flowering Plants: Three Conserved Lineages That Are at Least 200 Million Years Old and Possible Auto- and Cross-Regulation. Genetics 163, 311–319, doi:10.1093/genetics/163.1.311 (2003).

42 Zhang, H. et al. A Comprehensive Online Database for Exploring approximately 20,000 Public Arabidopsis RNA-Seq Libraries. Mol. Plant 13, 1231–1233, doi:10.1016/j.molp.2020.08.001 (2020).

43 Li, H. Minimap2: pairwise alignment for nucleotide sequences. Bioinformatics 34, 3094–3100, doi:10.1093/bioinformatics/bty191 (2018).

44 Emms, D. M. & Kelly, S. OrthoFinder: phylogenetic orthology inference for comparative genomics. Genome Biol. 20, 238, doi:10.1186/s13059-019-1832-y (2019).

